# Prenatal anti-oxidant treatment suppresses maternal immune activation induced increases in alcohol self-administration in a sex-specific manner

**DOI:** 10.1101/2025.06.16.660024

**Authors:** Skylar E. Nicholson, Kelly A. Hewitt, Cara S. Brauen, Angela M. Henricks

## Abstract

Prenatal exposure to infection is a risk factor in the development of several neuropsychiatric disorders, including schizophrenia and depression, which often co-occur with alcohol misuse, exacerbating adverse outcomes. Despite these consequences, little is understood regarding the mechanisms by which early exposure to infection might increase the risk of alcohol misuse. We have previously demonstrated that maternal immune activation (MIA) combined with adolescent alcohol exposure (AA) increases home-cage drinking and disrupts neural circuit function in adulthood. The current project aims to determine whether these findings extend to the motivation to work for alcohol, and whether prenatal anti-oxidant treatment can prevent the effects of MIA on self-administration. Pregnant Sprague-Dawley rats were exposed to poly(I:C) (4mg/kg) or saline on gestational day 15, and the anti-oxidant n-acetylcysteine (NAC; 100 mg/kg) or saline 24 hours before and after MIA. Offspring were given 24-hour home cage access to 10% ethanol and water dur adolescence. In adulthood, offspring were trained to self-administer 10% ethanol and tested on escalating schedules of reinforcement: fixed ratio (FR)1, FR2, FR4, and progressive ratios (PR). MIA and NAC independently led to an increased willingness to work for alcohol in males, compared to saline-treated rats. NAC treatment suppressed the effect of MIA on self-administration behavior. These data support the hypothesis that oxidative stress caused by MIA negatively influences development, priming the brain to be more susceptible to the negative effects of AA, leading to an increased risk of alcohol misuse in adulthood, particularly in males.

## 1. INTRODUCTION

Alcohol misuse and dependence are a source of substantial social, behavioral, economic, and health-related problems worldwide (Griswold et al., 2018). In the United States alone, chronic alcohol use accounts for approximately 54,000 deaths in men and 28,000 deaths in women annually (Esser, 2024). There is significant heterogeneity among individuals with alcohol use disorder (AUD) and the reasons they cite for developing problematic drinking (Witkiewitz et al., 2019), which suggests that alcohol misuse develops via multiple mechanisms and might require different therapies across different individuals (Litten et al., 2015).

Previous work indicates that vulnerability for problematic drinking likely stems from multiple factors in early life that alter the trajectory of brain development during critical periods (e.g., prenatal life and adolescence) (Guinle, 2020; Schepis et al., 2011). Such risk factors increase an individual’s susceptibility to stress and predisposition to neuropsychiatric disorders later in life. Prenatal exposure to infection is one of these risk factors, increasing the likelihood of the development of disorders like schizophrenia, bipolar disorders, and depression (Estes and McAllister, 2016), which all frequently co-occur with alcohol misuse (Grant et al., 2015). Similarly, adolescent stressors like drinking also increase the risk of developing alcohol misuse (Lees et al., 2020; Pfefferbaum et al., 2018), but no one stressor guarantees AUD development. The current study aims to begin investigating the mechanisms by which prenatal exposure to infection, combined with adolescent alcohol drinking (i.e., “dual-hits”), might increase the susceptibility for alcohol misuse later in life.

We previously showed that maternal immune activation (MIA), a well-validated rodent model of prenatal exposure to infection, combined with adolescent alcohol exposure (AA) increases home-cage alcohol intake in male and female adult offspring (Henricks et al., 2022). The MIA model results in a robust immune response in the pregnant rat, leading to cognitive, behavioral, and neurobiological pathologies that resemble mental illness (Estes and McAllister, 2016). In particular, MIA offspring exhibit increased pro-inflammatory cytokine secretion and microglia activation, abnormal synaptic plasticity, and deficits in social behaviors, exploratory behavior and sensorimotor gating (Cieślik et al., 2020; Meyer, 2014). AA is also associated with behavioral and neurobiological deficits, including neuroinflammation, synaptic plasticity changes, and executive functioning deficits (Lees et al., 2020; Spear, 2018). However, our data suggest neither MIA nor AA alone is sufficient to produce changes in adulthood alcohol drinking, but that these dual-hits lead to significant disruptions in the brain and behavior (Henricks et al., 2022).

The present study aims to better understand the mechanisms by which MIA might lead to an increased sensitivity to the negative effects of AA. Inflammation is closely related to oxidative stress, so it is not surprising that others have shown that MIA increases oxidative stress in the fetal brain (Pelição et al., 2016; Tapia-Rojas et al., 2019; Zawadzka et al., 2021; Zhang et al., 2021). Oxidative stress results from the overproduction of free radicals that is not controlled by anti-oxidant mechanisms, and appears to cause microglia activation and changes in synaptic function in the cortex (Simpson and Oliver, 2020; Tönnies and Trushina, 2017). Prenatal oxidative stress may therefore be a major mechanism by which MIA alters the trajectory of brain development and, subsequently, behavior in adulthood (Cieślik et al., 2020; Zawadzka et al., 2021). We therefore hypothesized that prenatal anti-oxidant treatment would prevent MIA+AA-induced increases in the motivation to consume alcohol in adulthood.

## 2. MATERIALS AND METHODS

### 2.1 Animals

All rats were housed in standard polycarbonate cages with *ad libitum* access to food and water. Rats were kept under a 12-hour reverse light-dark cycle (lights off at 08:00) and all experiments were conducted during the rats’ active phase to avoid circadian rhythm disruption. Animal facilities were accredited by the American Association of Laboratory Animal Care (AALAC) and all procedures were approved by the Institutional Animal Care and Use Committee (IACUC) at Washington State University.

### 2.2 Maternal immune activation and anti-oxidant treatment

Adult female Sprague-Daley rats (*n* = 13), obtained from Inotiv (Lafayette, IN, USA), underwent timed breeding. Female rats were continuously housed with male Sprague-Dawley rats and examined daily for a vaginal mucus plug, suggesting copulation. Day of confirmed copulation was considered gestational day (GD)0. On GD15, pregnant females received 4 mg/kg of polyinosinic:polycytidylic acid (poly(I:C); Tocris, Minneapolis, MN, USA) or saline via the tail vein. Blood was collected from each dam 3 hours after saline or poly(I:C) injection. Blood was centrifuged to acquire serum and stored at -20°C. We then measured pro-inflammatory cytokines IL6 and TNFα in dam serum using standard ELISA kits, following the manufacturer’s recommendations (RayBiotech, Peachtree Corners, GA, USA).

On GD14 and GD16, dams also received either saline or n-acetylcysteine (NAC; 100 mg/kg; Sigma Aldrich, St. Louis, MO, USA). NAC is a precursor of the anti-oxidant glutathione (Raghu et al., 2021), and reverses many of the cognitive and behavioral deficits induced by MIA (Lanté et al., 2008; Swanepoel et al., 2018). We ultimately had four experimental conditions: saline/saline (*n* = 9M, 9F from 3 dams), poly(I:C)/saline (*n* = 7M, 8F from 3 dams), saline/NAC (*n* = 9M, 9F from 3 dams), and poly(I:C)/NAC (*n* = 9M, 12F from 4 dams). Litters were culled to no more than 3 pups per sex to control for any litter effects, and pups were weaned on postnatal day (PD)21 and housed with same-sex litter mates.

### 2.3 Adolescent alcohol exposure

Pups were exposed to AA from PD28-48. Home cages were divided in half using perforated, plexiglass cage dividers, such that pups were separated to one pup per side, but still allowed some social contact with a conspecific. Each side of the cage included feeders with two bottles, one containing water and the other containing 10% ethanol (EtOH). The location of the water and EtOH bottles was counterbalanced. EtOH bottles and pups were weighed daily to track the g/kg of ethanol consumed during AA. On PD48, EtOH bottles and perforated cage dividers were removed, and pups were left to develop normally until PD70. In the current experiment, all pups received AA, since our previous research demonstrated that this adolescent exposure is necessary to unmask the effects of MIA on adulthood alcohol consumption (Henricks et al., 2022).

### 2.4 Alcohol self-administration training

Once the rats reached adulthood (PD70), they were trained to self-administer 10% EtOH in standard operant boxes (10” L x 12.5” W x 8.5” H) programmed with MED-Associates IV software. The operant boxes are equipped with two retractable levers on the wall of the chamber with a receptacle in the center for liquid distribution. Responses to the right lever—the active lever—resulted in the delivery of 0.1mL of liquid via a syringe pump. Responses to the left lever—the inactive lever—resulted in no liquid delivery. A house light located above the two levers was illuminated for the duration of each session.

Rats were trained to self-administer 10% EtOH using a sucrose-fading technique, in 30-minute sessions, 5-days a week (Monday – Friday) as described in our previous work (Henricks et al., 2016; Williams et al., 2012). Briefly, rats were trained to self-administering 10% sucrose for 4-5 sessions, then 10% sucrose and 10% EtOH for 5 sessions, then 5% sucrose and 10% EtOH for 5 sessions, then just 10% EtOH for 15 sessions under a fixed ratio-1 (FR1) schedule of reinforcement.

Following FR1 training, rats performed 3 additional self-administration sessions for each escalating schedule of reinforcement: FR2 (30 minutes), FR4 (30 minutes), and progressive ratio (PR; 60 minutes). This increasing schedule of reinforcement allowed us to measure the offspring’s motivation to work for alcohol.

### 2.5 Elevated Plus Maze

Since it has been demonstrated that MIA influences locomotor and anxiety-like behavior in adulthood (Henricks et al., 2022; Rahimi et al., 2020), which could impact self-administration behavior, we also measured Elevated Plus Maze (EPM) performance after self-administration was complete. Rats were placed in the center of the maze and allowed to freely explore the apparatus in dim lighting for 5 minutes. Sessions were recorded and the time spent in the closed and open arms, as well as total distance traveled, were analyzed using EthoVision XT (Noldus, Leesburg, VA). See Figure 1 for an experimental timeline.

**Figure 1:**
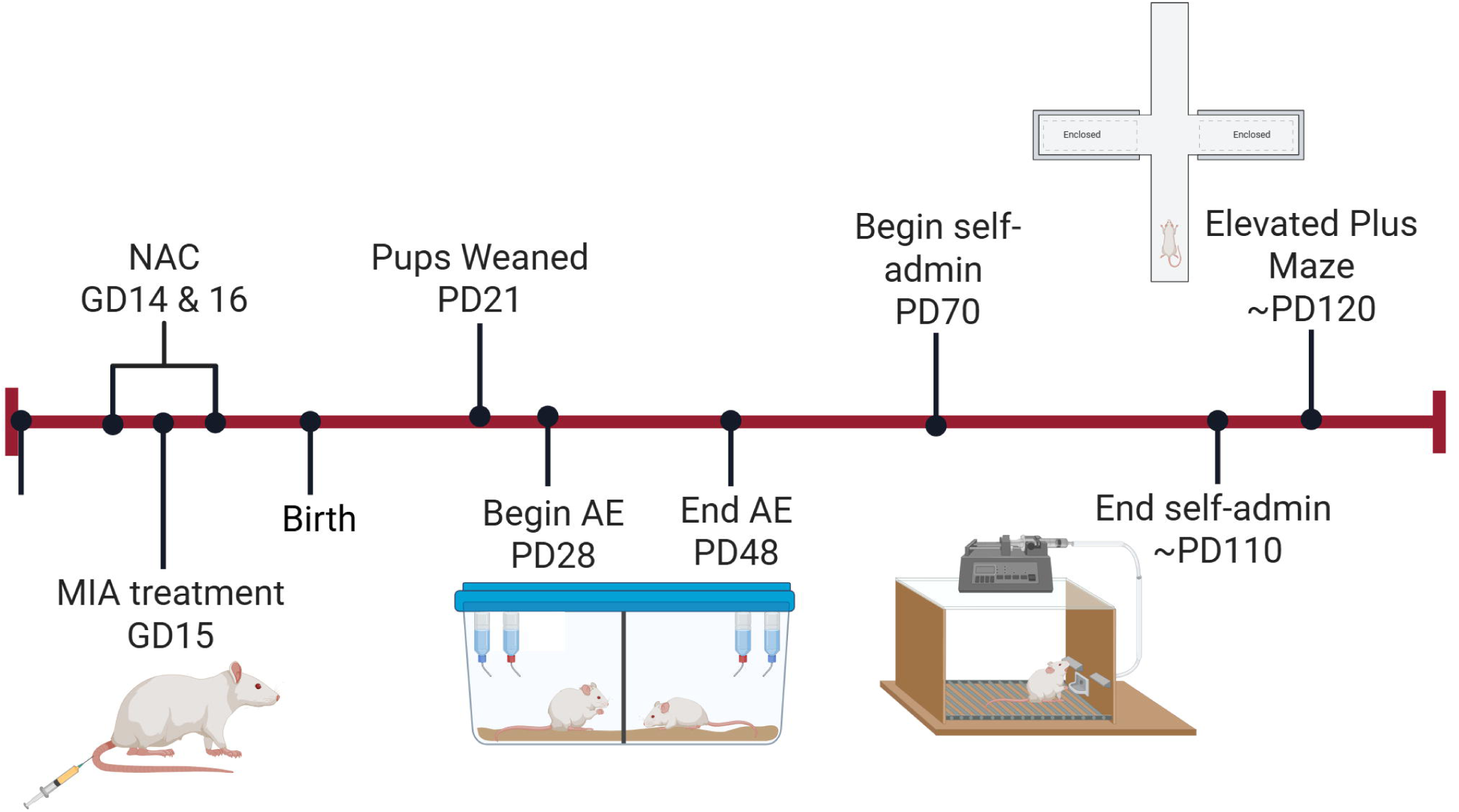
Experimental timeline.

## 3. STATISTICAL ANALYSES

### 3.1 Impact of Poly(I:C) and NAC on dams

Pro-inflammatory cytokine levels (IL6 and TNFα) and change in dam weight from GD14 to GD16 were analyzed using two-way ANOVAs with MIA (saline or poly(I:C)) and NAC treatment (saline or NAC) as the independent variables.

### 3.2 Alcohol drinking behavior

Average g/kg of alcohol consumed across the 20 days of AA was analyzed using a two-way ANOVA with MIA (saline or poly(I:C)) and NAC treatment (saline or NAC) as the independent variables. In adulthood, for the FR1 schedule of reinforcement, average rewards delivered and inactive lever presses over the last 5 sessions were analyzed using a two-way ANOVA with MIA (saline or poly(I:C)) and NAC treatment (saline or NAC) as the independent variables. A similar method was used to FR2, FR4 and PR, with the average rewards delivered or average breakpoint, as well as inactive lever presses within a self-administration session, serving as the dependent variables. Any interactions were followed up with Tukey’s *post-hoc* t-tests.

### 3.3 Elevated plus maze

Percent time spent in the closed and open arms, as well as total distance traveled were analyzed using a two-way ANOVA with MIA (saline or poly(I:C)) and NAC treatment (saline or NAC) as the independent variables.

To further control for litter effects, we used dam ID as a covariate in all offspring analyses. We also analyzed male and female data separately, since we had an *a priori* hypothesis that results would be sex-specific based on our and others’ previous work (Henricks et al., 2022; Liu et al., 2024).

## 4. RESULTS

### 4.1 Impact of Poly(I:C) and NAC on dams

For IL6, an ANOVA revealed no significant effects of MIA [*F*(1,9) = 0.19, *p* = 0.68, *n_p_^2^* = 0.02] or NAC [*F*(1,9) = 0.66, *p* = 0.44, *n_p_^2^* = 0.07], or a MIA*NAC interaction [*F*(1,9) = 0.72, *p* = 0.42, *n_p_^2^* = 0.07] (Figure 2A). For TNFα, an ANOVA revealed no significant effects of MIA [*F*(1,9) = 0.60, *p* = 0.46, *n_p_^2^* = 0.06] or NAC [*F*(1,9) = 0.20, *p* = 0.67, *n_p_^2^* = 0.02], or a MIA*NAC interaction [*F*(1,9) = 0.01, *p* = 0.91, *n_p_^2^* = 0.00] (Figure 2A). It is important to note that we previously demonstrated that this same dose of poly(I:C) increased IL6 and TNFα levels in dams two-hours after injection (Henricks et al., 2022), but we may be underpowered in the current experiment to detect any significant changes in cytokine levels.

**Figure 2:**
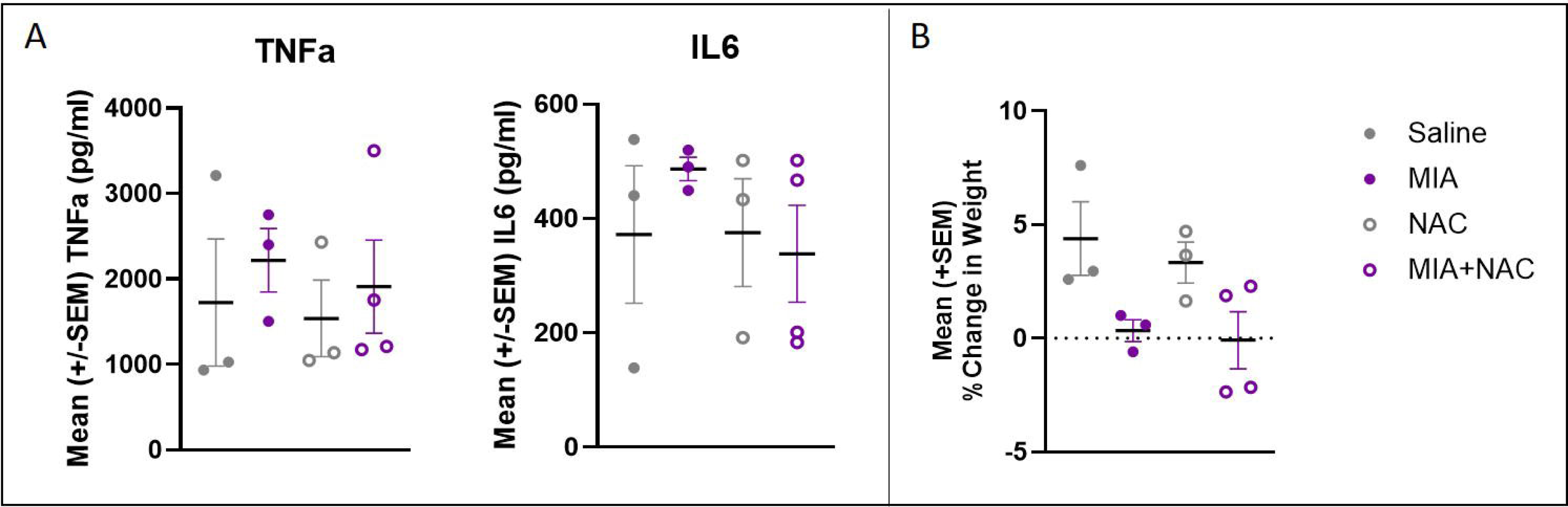
TNFα and IL6 serum levels in dams 3 hours after saline or poly(I:C) injection on GD15 (A). Percent change in weight from GD14 to GD16 in dams (B). There was a significant main effect of MIA, suggesting that the dams that received poly(I:C) gained less weight over that time period (*p* < 0.05).

An ANOVA revealed a significant effect of MIA [*F*(1,9) = 9.82, *p* < 0.05, *n_p_^2^* = 0.52] on weight change from GD14 to GD16, with dams that received poly(I:C) gaining less weight in that timeframe, similar to what we observed in previous studies (Henricks et al., 2022; Figure 2B). There was no significant effect of NAC [*F*(1,9) = 0.38, *p* = 0.55, *n_p_^2^* = 0.04] or a MIA*NAC interaction [*F*(1,9) = 0.07, *p* = 0.80, *n_p_^2^* = 0.01].

### 4.2 Adolescent alcohol drinking

In males and females, there was no effect of MIA [males: *F*(1,29) = 2.65, *p* = 0.11, *np^2^* = 0.08; females: *F*(1,29) = 2.89, *p* = 0.10, *np^2^* = 0.08] or NAC [males: *F*(1,29) = 0.37, *p* = 0.55, *np^2^* = 0.01; females: *F*(1,29) = 0.09, *p* = 0.76, *n_p_^2^* = 0.00], or an MIA*NAC interaction [males: *F*(1,29) = 3.20, *p* = 0.09, *n_p_^2^* = 0.10; females: *F*(1,29) = 1.70, *p* = 0.20, *n_p_^2^* = 0.05] for average g/kg of alcohol consumed in adolescence (Figure 3).

**Figure 3:**
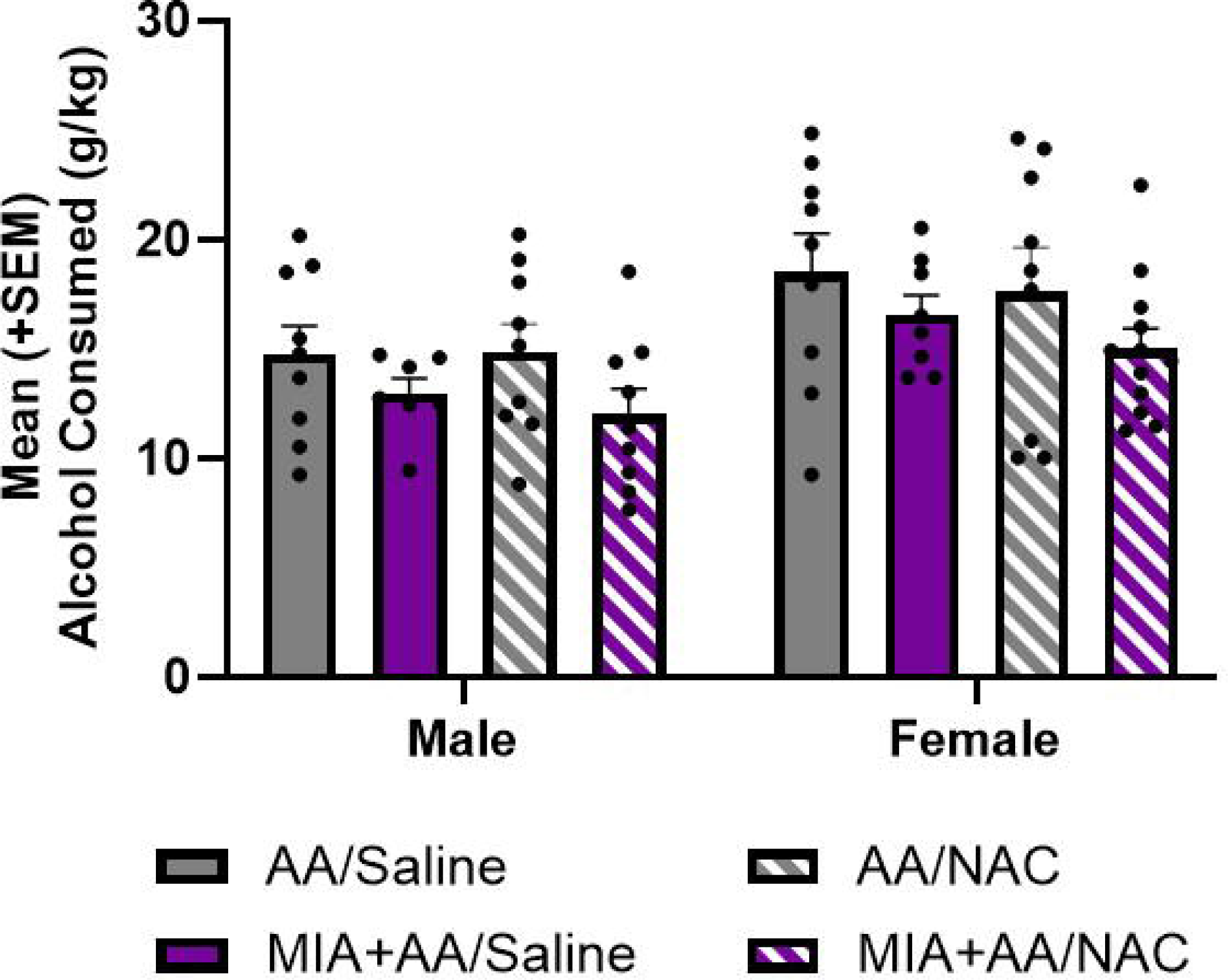
Average g/kg of alcohol consumed during adolescence. Neither MIA or NAC significantly influenced adolescent alcohol intake in either sex (*p*’s > 0.05; n = 7-12/group/sex).

### 4.3 Alcohol self-administration in adulthood

In males, for FR1, there was a significant effect of MIA [*F*(1,29) = 4.31, *p* < 0.05, *n_p_^2^*= 0.13], but no effect of NAC [*F*(1,29) = 0.23, *p* = 0.64, *n_p_^2^* = 0.01], and no MIA*NAC interaction [*F*(1,29) = 3.30, *p* = 0.08, *n_p_^2^* = 0.10] for number of awards administered (Figure 4A). For FR2, there was no significant effect of MIA [*F*(1,29) = 0.21, *p* = 0.65, *n_p_^2^* = 0.01], no effect of NAC [*F*(1,29) = 0.14, *p* = 0.71, *n_p_^2^* = 0.01], no MIA*NAC interaction [*F*(1,29) = 3.45, *p* = 0.07, *n_p_^2^* = 0.11] for number of rewards administered (Figure 4B). For FR4, there was no significant effect of MIA [*F*(1,29) = 0.52, *p* = 0.48, *n_p_^2^* = 0.02] and no effect of NAC [*F*(1,29) = 0.89, *p* = 0.35, *n_p_^2^* = 0.03] for number of rewards administered. However, there was a significant MIA*NAC interaction [*F*(1,29) = 6.12, *p* < 0.05, *n_p_^2^*= 0.17]. *Post-hoc* t-tests revealed that male offspring exposed to MIA and NAC, independently, received more rewards under an FR4 schedule of reinforcement compared to male offspring exposed to saline (*p’s* < 0.05) (Figure 4C). For PR, there was no significant effect of MIA [*F*(1,29) = 1.25, *p* = 0.27, *n_p_^2^* = 0.04] and no effect of NAC [*F*(1,29) = 2.44, *p* = 0.13, *n_p_^2^* = 0.08] on average break point. However, there was a significant MIA*NAC interaction [*F*(1,29) = 9.56, *p* < 0.01, *n_p_^2^* = 0.25]. *Post-hoc* t-tests revealed that male offspring exposed to MIA and NAC, independently, had higher PR breakpoints compared to male offspring exposed to saline (*p’s* < 0.01) (Figure 4D).

**Figure 4:**
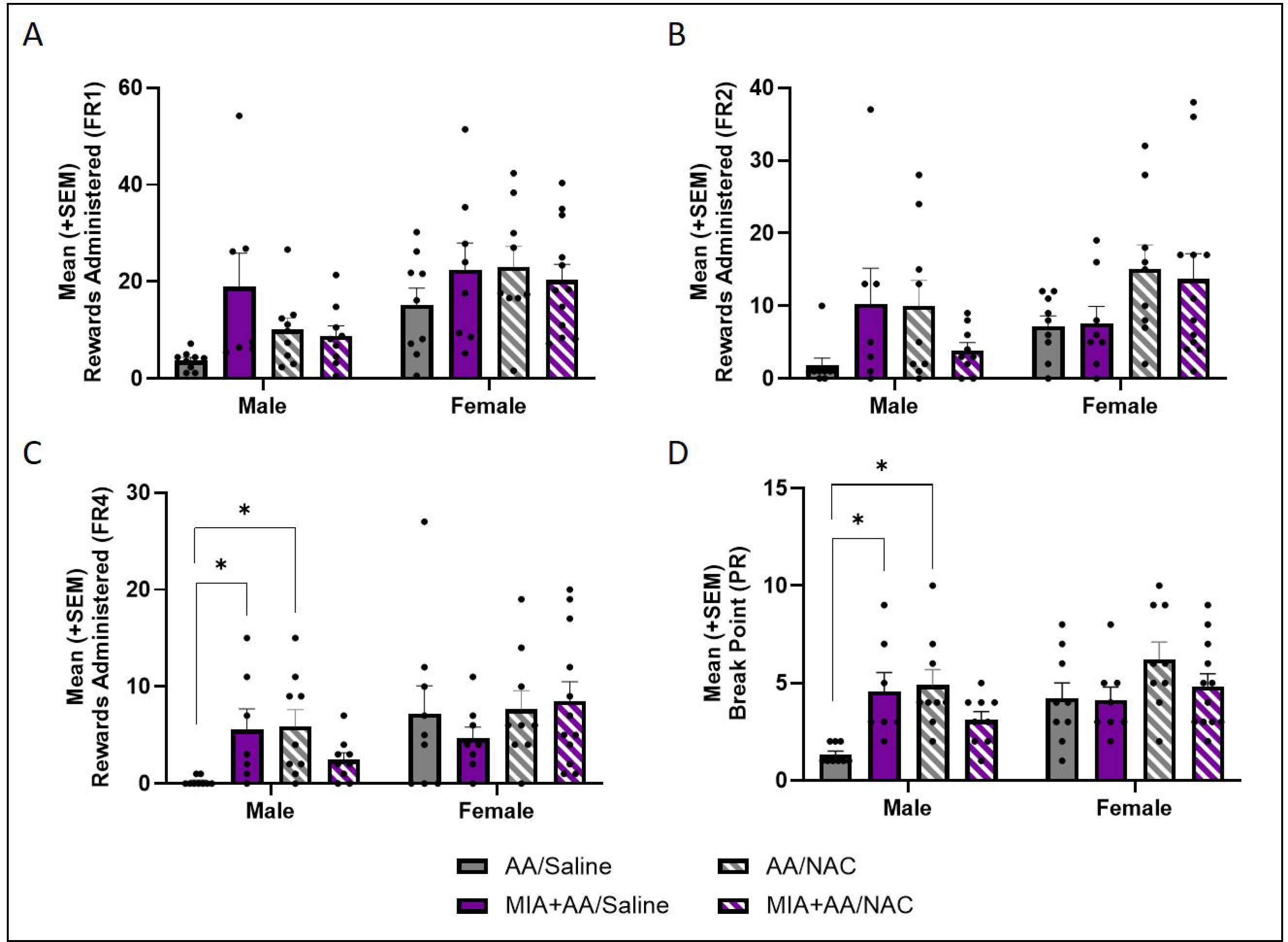
Average rewards administered in each sex during FR1 (A), FR2 (B) and FR4 (C) schedules of reinforcement, as well as average break point for the PR schedule (D). In males, both MIA and NAC independently led to increased rewards administered in FR4 and break point in PR compared to saline-treated offspring (\**p*’s < 0.05; n = 7-12/group/sex).

In females, for FR1, there was no significant effect of MIA [*F*(1,33) = 0.29, *p* = 0.59, *n_p_^2^* = 0.01] and no effect of NAC [*F*(1,33) = 0.10, *p* = 0.76, *n_p_^2^* = 0.00] for number or rewards administered. There was a significant MIA*NAC interaction [*F*(1,33) = 4.30, *p* < 0.05, *n_p_^2^* = 0.12], but *post-hoc* t-tests revealed no significant differences between individual groups (*p’s* > 0.05) (Figure 4A). For FR2, there was no significant effect of MIA [*F*(1,33) = 0.05, *p* = 0.83, *np^2^* = 0.00], no effect of NAC [*F*(1,33) = 3.57, *p* = 0.07, *n_p_^2^* = 0.10], and no MIA*NAC interaction [*F*(1,33) = 2.58, *p* = 0.12, *n_p_^2^* = 0.07] for number of rewards administered (Figure 4B). For FR4, there was no significant effect of MIA [*F*(1,33) = 0.18, *p* = 0.67, *n_p_^2^* = 0.01], no effect of NAC [*F*(1,33) = 0.52, *p* = 0.48, *n_p_^2^* = 0.02], no MIA*NAC interaction [*F*(1,33) = 0.00, *p* = 0.99, *n_p_^2^* = 0.00] for number of rewards administered (Figure 4C). For PR, there was no significant effect of MIA [*F*(1,33) = 1.21, *p* = 0.28, *n_p_^2^* = 0.04] and no effect of NAC [*F*(1,33) = 1.80, *p* = 0.19, *n_p_^2^* = 0.05] for average break point. However, there was a significant MIA*NAC interaction [*F*(1,33) = 4.74, *p* < 0.05, *n_p_^2^*= 0.13], but *post-hoc* t-tests revealed no significant differences between individual groups (*p’s* > 0.05) (Figure 4D). Inactive lever presses were not significantly altered by MIA or NAC in either sex (see supplementary materials).

### 4.4 Elevated plus maze

In males, there was a significant effect of MIA [*F*(1,29) = 7.04, *p* < 0.05, *n_p_^2^* = 0.20] and NAC [*F*(1,29) = 5.61, *p* < 0.05, *n_p_^2^* = 0.16] on total distance traveled, but no MIA*NAC interaction [*F*(1,29) = 0.01, *p* = 0.93, *n_p_^2^* = 0.00] (Figure 5A). There was no effect of MIA [*F*(1,29) = 0.39, *p* = 0.54, *n_p_^2^* = 0.01], NAC [*F*(1,29) = 0.26, *p* = 0.62, *n_p_^2^* = 0.01], or a NAC*MIA interaction [*F*(1,29) = 1.07, *p* = 0.31, *n_p_^2^* = 0.04] for percent time spent in the closed arms (Figure 5B). Similarly, there was no effect of MIA [*F*(1,29) = 0.00, *p* = 1.00, *n_p_^2^* = 0.00], NAC [*F*(1,29) = 0.82, *p* = 0.37, *n_p_^2^* = 0.03], or a NAC*MIA interaction [*F*(1,29) = 0.61, *p* = 0.44, *n_p_^2^* = 0.02] for percent time spent in the open arms (Figure 5C).

**Figure 5:**
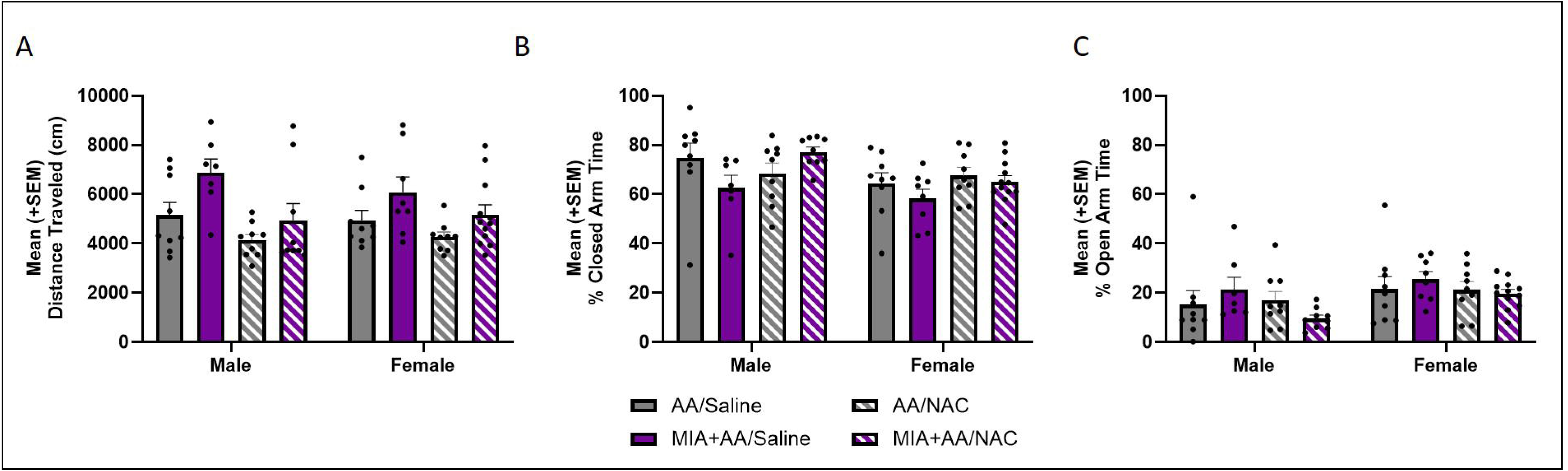
Average distance traveled (A) and percent time spent in the closed (B) and open arms (C) of the elevated plus maze. There was a significant main effect of MIA and NAC on distance traveled in males, and a main effect of MIA on distance traveled in females (*p*’s < 0.05). MIA was associated with increased distance traveled in both sexes, while NAC was associated with reduced distance traveled in males (n = 7-12/group/sex).

In females, there was a significant effect of MIA [*F*(1,33) = 5.68, *p* < 0.05, *np^2^* = 0.15], but no effect of NAC [*F*(1,33) = 2.76, *p* = 0.11, *np^2^* = 0.08] or an MIA*NAC interaction [*F*(1,33) = 0.02, *p* = 0.88, *np^2^* = 0.00] on total distance traveled (Figure 5A). There was no effect of MIA [*F*(1,33) = 1.82, *p* = 0.19, *np^2^* = 0.05], NAC [*F*(1,33) = 1.02, *p* = 0.32, *np^2^* = 0.03], or a NAC*MIA interaction [*F*(1,33) = 0.49, *p* = 0.49, *np^2^* = 0.02] for percent time spent in the closed arms (Figure 5B). There also was no effect of MIA [*F*(1,33) = 0.15, *p* = 0.70, *np^2^* = 0.01], NAC [*F*(1,33) = 0.32, *p* = 0.58, *n_p_^2^* = 0.01], or a NAC*MIA interaction [*F*(1,33) = 0.04, *p* = 0.85, *n_p_^2^* = 0.00] for percent time spent in the open arms (Figure 5C).

## 5. DISCUSSION

The current experiment demonstrates that MIA+AA induces sex-specific increases in the motivation to drink alcohol in adulthood. Male MIA+AA rats were willing to work harder for alcohol based on increased responding on FR4 and PR schedules compared to males exposed to AA alone. Interestingly, we found similar effects in male NAC+AA rats (those not exposed to MIA), but NAC also suppressed the effect of MIA+AA on motivation to drink alcohol. However, none of these affects were apparent in the female rats. Further, neither MIA nor NAC impacted overall adolescent alcohol intake in either sex. Our findings therefore suggest that males are more sensitive to the effects of MIA+AA on alcohol self-administration, and that prenatal anti-oxidant treatment can attenuate these effects.

These data support other studies showing sex-specific effects of MIA. For example, male MIA offspring show more spatial working memory, motor and social interaction deficits compared to MIA females (Gogos et al., 2020; Haida et al., 2019; Liu et al., 2024; Xuan and Hampson, 2014). Sex hormones may contribute to these effects since estrogens enhance immunocompetence, whereas testosterone inhibits immunocompetence (Roved et al., 2017). It has therefore been hypothesized that males have a higher inflammatory tone than females (McCarthy, 2019), which could explain the increased susceptibility of male rats to the adverse effects of MIA. It is important to note, however, that females are underrepresented in MIA research (Coiro and Pollak, 2019), so our current understanding of how MIA might impact males and females differently is in its infancy, and future research should not only represent both sexes, but should also include hormone manipulations in order to better understand the mechanisms driving any sex-specific effects.

As we hypothesized, prenatal anti-oxidant treatment with NAC suppressed the effect of MIA+AA on motivation to drink alcohol in adult males. These data support our hypothesis that oxidative stress is an important contributor to the effects of MIA in adulthood. Prenatal oxidative stress has been implicated as a risk factor for several psychiatric disorders (al-Haddad et al., 2019; Usui et al., 2023; Xu et al., 2019), perhaps due to the impact of oxidative stress on neurodevelopment. Oxidative balance is vital for processes like neurogenesis and synaptic pruning (Londono Tobon et al., 2016), so disruptions in this system via MIA might make organisms more susceptible to other environmental risk factors during critical periods of development, like adolescence. There is evidence for this hypothesis, as MIA leads to oxidative stress-dependent reductions in synaptic plasticity in the hippocampus in adolescent male offspring (Lanté et al., 2007). Our future work therefore aims to dissect the mechanisms by which MIA-induced oxidative stress primes the brain to be susceptible to second “hits,” like AA, leading to maladaptive neural circuit changes in adulthood that are associated with alcohol misuse.

A particularly interesting and novel finding of this study is that prenatal NAC alone led to increased motivation to consume alcohol in adult males. NAC stimulates the production of glutathione, which the body produces from various amino acids to reduce reactive oxygen species (ROS) (Ballatori et al., 2009; Detcheverry et al., 2023; Townsend et al., 2003), so in the presence of oxidative stress by MIA, glutathione production is beneficial. However, when MIA-induced ROS are not present, NAC administration might disrupt glutathione homeostasis, causing detrimental effects. For instance, others have shown that increased glutathione is associated with the development of tumors and resistance to chemotherapy (Kennedy et al., 2020). In our study, it appears that too much glutathione might disrupt the oxidative balance in cells and ultimately contribute to increased motivation to consume alcohol after AA. This finding limits the applicability of NAC as a preemptive treatment for MIA, but the ability of NAC to attenuate effects of MIA+AA confirm that oxidative stress is an important mechanism by which prenatal exposure to infection produces long-term effects, and requires further investigation.

The EPM data indicate that MIA influences locomotor behavior in both sexes, and that NAC alone influences locomotor behavior in males, though these effects were independent since there was no MIA*NAC interaction. While MIA has been shown to cause an increase, decrease, or no change in locomotor activity (Henricks et al., 2022; Tellez-Merlo et al., 2019; Vorhees et al., 2015), in the present study MIA appeared to increase overall distance traveled in the EPM. Locomotor changes could have contributed to the increase in active lever presses observed in male MIA+AA offspring. However, inactive lever presses were not influenced by MIA or NAC in males in this study, suggesting that any increase in motivation to consume alcohol is not simply reflective of a change in locomotor behavior.

One limitation of this study is that all rats were exposed to AA, so we did not have a pure control group. However, our primary purpose was to test whether oxidative stress is a major mechanism by which MIA disrupts behavior, and since our previous work showed that AA was necessary to unmask the effects of MIA on adulthood drinking, we chose to minimize our animal use to the most relevant experimental groups. That being said, considering that NAC+AA impacted drinking in this study, it would be important in the future to determine whether prenatal NAC exposure alone (without AA) also leads to increased drinking in adulthood. Second, serum IL6 and TNFα were not significantly elevated by MIA in the current experiment. We largely contribute this to a lack of power with only 3-4 dams per experimental group since our previous work showed that the same dose of poly(I:C) increases IL6 and TNFα with a larger sample size (Henricks et al., 2022). Further, since we were able to replicate our and others previous findings that poly(I:C) prevents adequate weight gain in dams for the first 24-48 hours after treatment (Henricks et al., 2022; Howland et al., 2012), we feel confident that our protocol induced MIA in dams. That being said, our future work aims to determine whether there are more reliable biomarkers for inflammation after poly(I:C) treatment.

Overall, the current data significantly contribute to our understanding of the mechanisms by which MIA leads to long-term changes in alcohol drinking behaviors, and our future work aims to understand how MIA-induced oxidative stress and AA interact to disrupt neurodevelopment. Since others have shown that both MIA alone and AA alone reduce cortical dendritic spine density (Amodeo et al., 2021; Coiro and Pollak, 2019; Jury et al., 2017; Pekala et al., 2021), we specifically aim to determine the impact of MIA on cortical synaptic plasticity in adolescence, and whether oxidative stress drives these effects. Our ultimate, long-term goal is to identify how early life environmental stressors change brain development in such a way as to predispose individuals to alcohol misuse and improve translatability of preclinical findings by identifying biomarkers for future therapeutic development.

### Author credit

*Nicholson, Skylar*: Methodology, formal analysis, investigation, writing – original draft, visualization; *Hewitt, Kelly*: Formal analysis, investigation, writing – review & editing; *Brauen, Cara*: Investigation; *Henricks, Angela*: Conceptualization, methodology, formal analysis, resources, writing – review & editing, visualization, supervision, funding acquisition.

### Funding sources

This work was supported by a WSU Alcohol and Drug Abuse Research Program (ADARP) pilot grant, and the WSU Department of Psychology.

