## Supplementary figures and images for "Prenatal anti-oxidant treatment suppresses maternal immune activation induced increases in alcohol self-administration in a sex-specific manner"

### Supp Figure 1

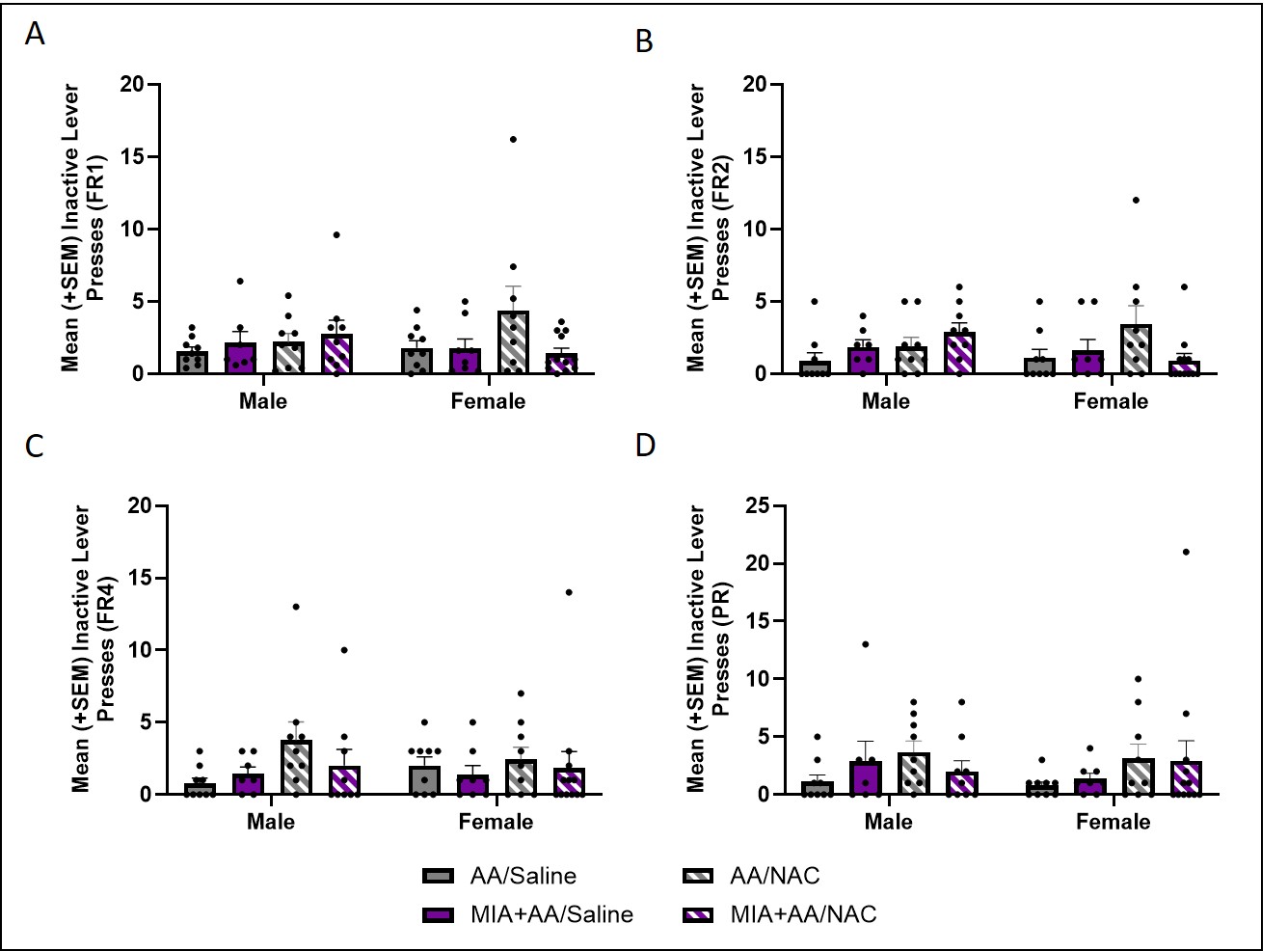
