## Supplementary material for "Prenatal anti-oxidant treatment suppresses maternal immune activation induced increases in alcohol self-administration in a sex-specific manner": Supp materials

Assistant Professor

Department of Psychology

Washington State University

Pullman, WA 99163

**Results**

*Inactive lever presses*

In males, for FR1, there no effect of MIA [*F*(1,29) = 1.45, *p* = 0.24, *n_p_^2^* = 0.05], NAC [*F*(1,29) = 2.16, *p* = 0.15, *n_p_^2^* = 0.07] or a MIA*NAC interaction [*F*(1,29) = 1.42, *p* = 0.24, *n_p_^2^* = 0.05] for average inactive lever presses (Supplemental Figure 1A). Similarly, there was no effect of MIA, NAC or a MIA*NAC interaction for inactive lever presses in FR2 (MIA: [*F*(1,29) = 2.55, *p* = 0.12, *n_p_^2^* = 0.08]; NAC: [*F*(1,29) = 2.62, *p* = 0.12, *n_p_^2^* = 0.08]; MIA*NAC: [*F*(1,29) = 0.01, *p* = 0.94, *n_p_^2^* = 0.00]), FR4 (MIA: [*F*(1,29) = 0.23, *p* = 0.64, *n_p_^2^* = 0.01]; NAC: [*F*(1,29) = 3.83, *p* = 0.06, *n_p_^2^* = 0.12]; MIA*NAC: [*F*(1,29) = 0.45, *p* = 0.51, *n_p_^2^* = 0.02]), and PR schedules of reinforcement (MIA: [*F*(1,29) = 0.04, *p* = 0.84, *n_p_^2^* = 0.00]; NAC: [*F*(1,29) = 1.07, *p* = 0.31, *n_p_^2^* = 0.04]; MIA*NAC: [*F*(1,29) = 0.49, *p* = 0.49, *n_p_^2^* = 0.02]) (Supplemental Figure 1B-D).

In females, there no effect of MIA [*F*(1,33) = 2.70, *p* = 0.11, *n_p_^2^* = 0.08], NAC [*F*(1,33) = 1.22, *p* = 0.28, *n_p_^2^* = 0.04] or a MIA*NAC interaction [*F*(1,33) = 2.31, *p* = 0.14, *n_p_^2^* = 0.07] for average inactive lever presses (Supplemental Figure 1A). Similarly, there was no effect of MIA, NAC or a MIA*NAC interaction for inactive lever presses in FR4 (MIA: [*F*(1,33) = 0.43, *p* = 0.52, *n_p_^2^* = 0.01]; NAC: [*F*(1,33) = 0.47, *p* = 0.50, *n_p_^2^* = 0.01]; MIA*NAC: [*F*(1,33) = 0.28, *p* = 0.60, *n_p_^2^* = 0.01] and PR schedules of reinforcement (MIA: [*F*(1,33) = 0.03, *p* = 0.87, *n_p_^2^* = 0.00]; NAC: [*F*(1,33) = 2.76, *p* = 0.11, *n_p_^2^* = 0.08]; MIA*NAC: [*F*(1,33) = 0.08, *p* = 0.79, *n_p_^2^* = 0.00]; (Supplemental Figure 1C&D). However, while there was no main effect of MIA [*F*(1,33) = 1.62, *p* = 0.21, *n_p_^2^* = 0.05] or NAC [*F*(1,33) = 0.53, *p* = 0.47, *n_p_^2^* = 0.02], there was a significant MIA*NAC interaction [*F*(1,33) = 5.13, *p* < 0.05, *n_p_^2^* = 0.14] for FR2 inactive lever presses (Supplemental Figure 1B), but *post-hoc* t-tests revealed no significant differences between individual groups (*p’s* > 0.05).

**Figure captions**

**Supplemental Figure 1:** Average inactive lever presses for FR1 (A), FR2 (B), FR4 (C), and PR (D) schedules of reinforcement (n=7-12/group/sex).
